# HKDC1 Promotes Liver Cancer Stemness Under Hypoxia via Stabilizing β-Catenin

**DOI:** 10.1101/2024.02.27.581958

**Authors:** Li Fan, Cheng Tian, Wentao Yang, Xiaoli Liu, Yogesh Dhungana, Haiyan Tan, Evan S Glazer, Jiyang Yu, Junmin Peng, Lichun Ma, Min Ni, Liqin Zhu

## Abstract

**Background and Aims:** Hexokinases (HKs), a group of enzymes catalyzing the first step of glycolysis, have been shown to play important roles in liver metabolism and tumorigenesis. Our recent studies identified HKDC1 as a top candidate associated with liver cancer metastasis. We aimed to compare its cell-type specificity with other HKs upregulated in liver cancer and investigate the molecular mechanisms underlying its involvement in liver cancer metastasis.

**Approach and Results:** We found that, compared to HK1 and HK2, the other two commonly upregulated HKs in liver cancer, HKDC1 was most strongly associated with the metastasis potential of tumors and organoids derived from two liver cancer mouse models we previously established. RNA in situ hybridization and single-cell RNA-seq analysis revealed that HKDC1 was specifically upregulated in malignant cells in hepatocellular carcinoma (HCC) and cholangiocarcinoma (CCA) patient tumors, whereas HK1 and HK2 were widespread across various tumor microenvironment lineages. An unbiased metabolomic profiling demonstrated that HKDC1 overexpression in HCC cells led to metabolic alterations distinct from those from HK1 and HK2 overexpression, with HKDC1 particularly impacting the tricarboxylic acid (TCA) cycle. HKDC1 was prometastatic in HCC orthotopic and tail vein injection mouse models and, molecularly, HKDC1 was induced by hypoxia and bound to glycogen synthase kinase 3β to stabilize β-catenin, leading to enhanced stemness of HCC cells.

**Conclusions:** Overall, our findings underscore HKDC1 as a prometastatic HK specifically expressed in the malignant compartment of primary liver tumors, thereby providing a mechanistic basis for targeting this enzyme in advanced liver cancer.

## INTRODUCTION

Hexokinases (HKs) are a family of enzymes that catalyzes the first step in glucose metabolism, the phosphorylation of glucose into glucose 6-phosphate (G6P) (1). There are four conventional HK isoforms, HK1, HK2, HK3 and HK4. They are similar in structure but different in their enzymatic efficacy and tissue specificity (2). The liver, one of the most important metabolic organs of the body, predominantly uses HK4 (also known as glucokinase, or GCK) to phosphorylate glucose. In hepatocellular carcinoma (HCC), HK4 is downregulated and HK1 and HK2 are upregulated to support tumorigenesis (3-5). Hexokinase domain containing 1 (HKDC1) is a fifth human hexokinase discovered recently via its association with gestational diabetes (6-8). HKDC1 has a low hexokinase activity compared to the other HKs and is nearly undetectable in the normal liver tissue (9, 10). Subsequent studies found that HKDC1 is significantly upregulated and is a protumorigenic player in HCC and other cancers (11-14). Its impact on glucose update of cancer cells and the subcellular association with mitochondria-associated membranes have also been demonstrated (15). What remains unexplored are the potential differences of HKDC1 compared to the other upregulated HKs regarding their lineage specificity and metabolic activity in HCC since HCC is well known for its complex tumor microenvironment (TME) which consists of various cell types with different functions and metabolic needs.

In this study, our group independently identified *Hkdc1* as one of the top upregulated genes in the metastatic tumors and organoids derived from an HCC genetic mouse model we previously established. Tumorigenesis in this genetic mouse model is driven by the dual loss of *Pten* and *Tp53* tumor suppressor genes in a *Prom1*-expressing liver progenitor population (*Prom1^CreERT2^; Pten^flx/flx^; Tp53^flx/flx^; Rosa-ZsGreen*, or PPTR) (16). Orthotopic tumors developed from the tumor organoids generated from this model gradually acquired cellular and molecular features of cholangiocarcinoma (CCA), the second most common primary liver cancer, and showed a significantly increased metastatic potential (17). We followed up these findings in this study and compared the distribution of HKDC1 to that of HK1 and HK2 in HCC and CCA patient tumors. For functional and metabolic analyses, we mainly used human hepatocellular cancer cell lines and orthotopic xenograft models given the rarity of human CCA cell lines. Because of its upregulation in PPTR tumor organoids that exhibited strong liver cancer stem cell traits (17), we directed our focus towards understanding the involvement of HKDC1 in hypoxia and cancer stem cells and examined its participation in WNT/β-catenin signaling for the known importance of this pathway in supporting the stemness of both normal and malignant liver cells (18, 19).

## RESULTS

### HKDC1 is a top gene upregulated in metastatic tumors and organoids in PPTR liver cancer mouse model

We previously generated a liver cancer mouse model by targeting a *Prom1*-expressing liver progenitor population with the dual loss of tumor suppressor genes *Pten* and *Tp53*, (*Prominin1^CreERT2^; Pten^flx/flx^; Tp53^flx/flx^; Rosa^ZsG^*, or PPTR) (16). PPTR mice developed frequent primary tumors in the liver with histological features of HCC. But metastases were rare in this model. We then generated multiple cancer organoid lines derived from the PPTR primary tumors and they showed varing levels of tumorigenicity when orthotopically transplanted into the mouse liver. A subset of PPTR tumor organoids demonstrated a high frequency of metastatic tumor formation in the orthotopic allograft model at high frequencies, while others failed to grow in vivo (**Figure 1A**) (17). Tumors in the PPTR allograft model acquired CCA features, potentially due to a hepatocyte-to-cholangiocyte trans-differentiation (17). RNA-seq transcriptomic profiling of the PPTR genetic and allograft tumors as well as the tumor organoids was previously performed (GSE94583). In this study, comparative analyses were conducted to identify genes associated with enhanced metastasis in the allograft model. Similar analyses were performed for a hepatoblastoma mouse model we established by engineering an activating *Notch* mutation, *NICD* (<u>N</u>otch inter<u>c</u>ellular <u>d</u>omain), into the *Prom1*-expressing liver progenitors (*Prominin1^CreERT2^; Rosa^NICD1/+^; Rosa^ZsG^*, or PNR). Combining analyses from both mouse models enabled us to identify genes commonly associated with liver cancer metastasis.

**Figure 1.**
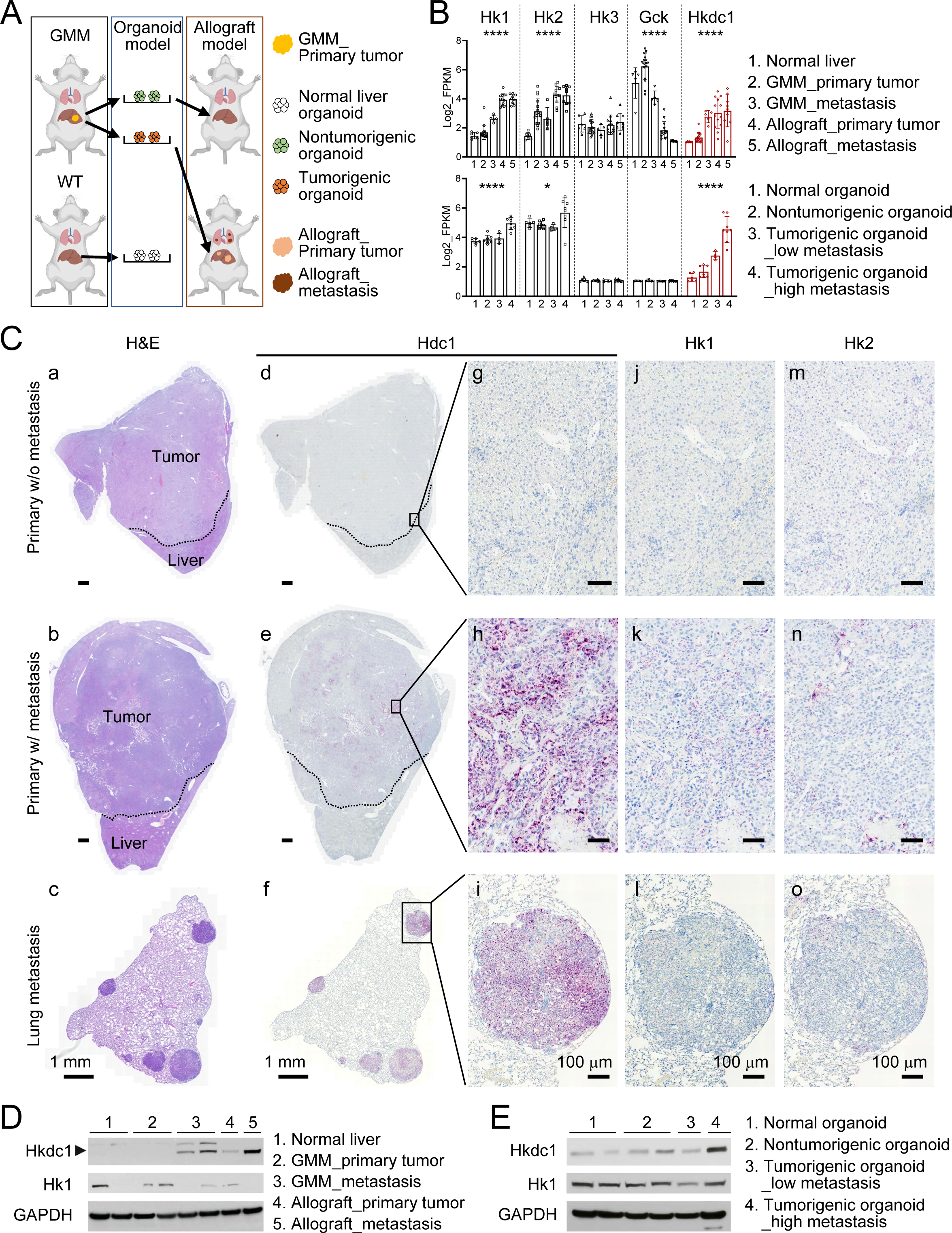
HKDC1 is upregulated in metastatic tumors and organoids in PPTR liver cancer mouse model. **A** A schemtic illustration for the establishment of PPTR genetic mouse model, organoid model, and orthotopic allograft models (generated by BioRender). **B** Quantitative comparison of the gene expression of the five HKs in PPTR tumor tissues (n=6, 16, 5, 13, and 9 for the five indicated groups, respectively) and organoids (n=6, 7, 4, and 8 for the four indicated groups, respectively). One-way ANOVA test: *P* values: * < 0.05; *** < 0.0001. **C** Hematoxylin and eosin (H&E) (a-c) and RNAscope staining of *Hkdc1* (d-i), *Hk1* (j-l), and *Hk2* (m-o) on the serial sections from the indicated tumors in the PPTR genetic mouse model. Images on the same column share the same scale bar. **D** Immunoblotting of Hkdc1 and Hk1 in the normal liver and indicated PPTR tumors. **E** Immunoblotting of Hkdc1 and Hk1 in the normal liver organoids and indicated PPTR tumor organoids.

We found that *Hkdc1*, a recently identified HK family member, was among the top genes significantly upregulated in the metastatic PNR and PPTR tumors and organoids (**Figure 1B** and **Supplemental Figure 1)**. Although the other two HKs commonly upregulated in liver cancer, *Hk1* and *Hk2*, exhibited similar upregulation in the metastatic PPTR and PNR tumors, their association with the metastatic potential of the tumor organoid was less pronounced compared to *Hkdc1*. To validate these findings, we performed RNAscope in situ hybridization using primary and metastatic tumors from the PPTR genetic mouse model. *Hkdc1* mRNA was detected in the metastatic tumor at a much higher level than that of *Hk1* and *Hk2* (**Figure 1C**). Immunohistochemistry (IHC) of Hkdc1 was not successful due to the lack of a mouse Hkdc1 antibody suitable for histological examination. Since Hkdc1 is structurally similar to HK1, we further compared their protein levels in the PPTR tumors and organoids via immunoblotting. We confirmed the increased levels of Hkdc1 protein in PPTR metastatic tumors (**Figure 1D**) and metastatic organoids (**Figure 1E**) but not that of HK1. Overall, among the HK family members, Hkdc1 showed the strongest association with the metastasis in the PPTR mouse model.

### HKDC1 upregulation is highly specific to malignant cells in HCC and CCA patients

Expression of HKDC1 in human HCC and CCA has been previously reported (18-20). However, liver cancer is known for its complex tumor microenvironment (TME) with the presence of various non-malignant cell populations in the tumor. The cell-type specificity of HKDC1 and other HKs have not been characterized, which could suggest their potentially different contributions to liver tumorigenesis. Thus, we examined the expression of *HK1*, *HK2*, and *HKDC1* in malignant cells and non-malignant cells using a previously reported single-cell RNA sequencing (scRNA-seq) of 46 HCC and CCA tumors (20). We found that *HKDC1* expression was highly specific to the malignant cells, whereas *HK1* and *HK2* expression was detected in both malignant and non-malignant compartments (**Figure 2A**). Consistently, t-SNE analysis of all the 17,164 malignant cells and 35,625 non-malignant cells combined from both cancer types in this dataset showed that the majority *HKDC1*^+^ cells were malignant while *HK1* and *HK2* expression was predominantly found in non-malignant cells (**Figure 2B**). To assess the expression of these genes in normal tissue and tumor, we further analyzed a multiregional scRNA-seq dataset generated from seven HCC and CCA patient tumors and their adjacent normal liver tissues (21). *HKDC1* expression was detected mostly in cholangiocytes in the tumor-adjacent liver, malignant tumor cells and some T cells in the tumor tissues (**Figure 2C**). In Contrast, *HK1* and *HK2* both showed a wide range of expression across nearly all cell types within the HCC and CCA TME (**Figure 2C**). *HK1* levels were particularly high in tumor-associated macrophages (TAMs) and cancer-associated fibroblasts (CAFs) (**Figure 2C**). In non-tumor tissues, HK1 was mainly expressed in the cholangiocytes rather than hepatocytes, while HK2 levels were very low in both cell types (**Figure 2C**).

**Figure 2.**
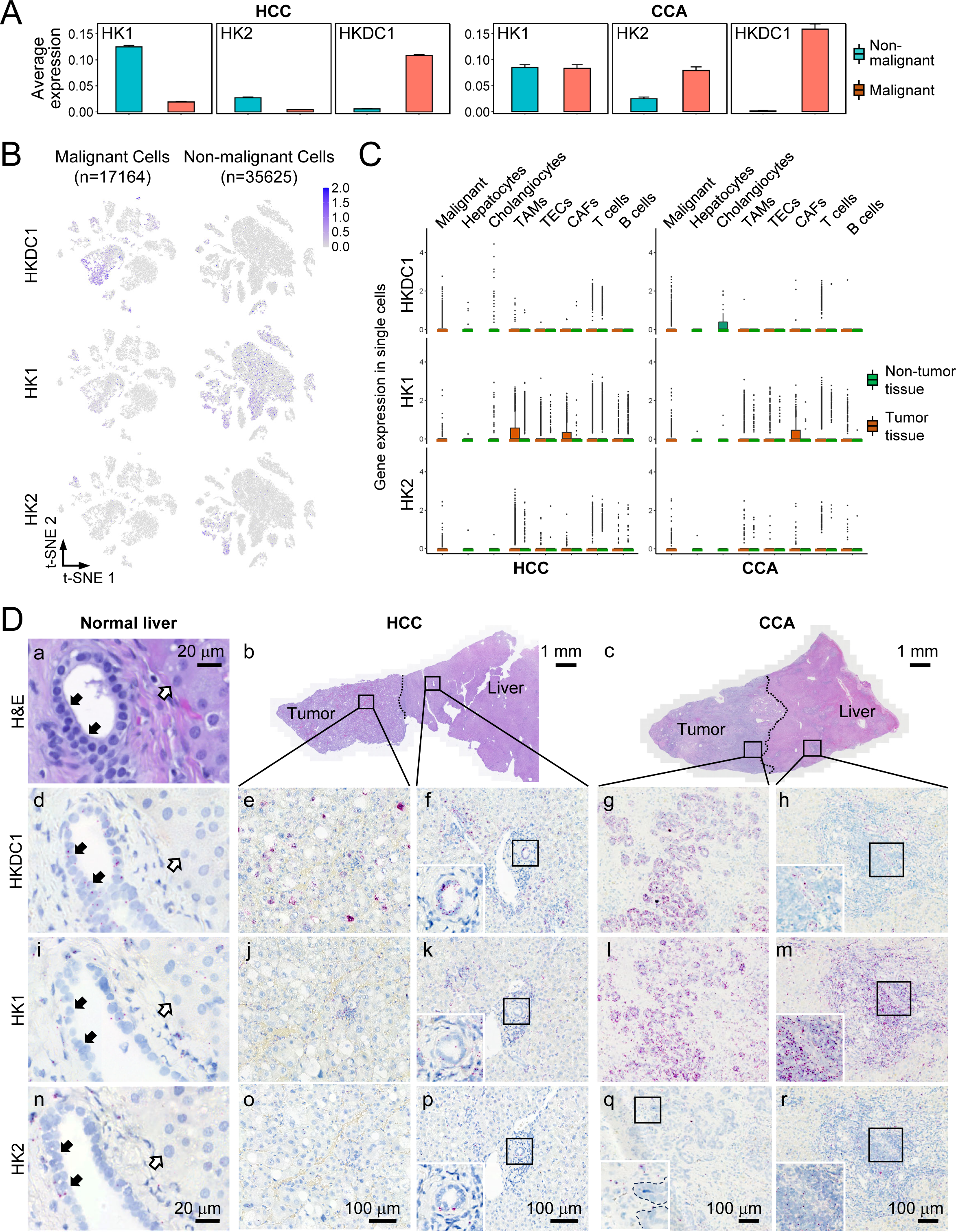
HKDC1 upregulation is highly specific to malignant cells in HCC and CCA patients. **A** Mean expression of *HK1*, *HK2*, and *HKDC1* in non-malignant cells and malignant cells from 46 published liver cancer patient samples including 25 HCC (left panel) and 12 CCA (right panel) patients (20). **B** tSNE plots of *HKDC1*, *HK1*, and *HK2* expression in the malignant and non-malignant cells in the samples in **A**. **C** *HKDC1*, *HK1*, and *HK2* expression in individual cell types using the published scRNA-seq data from 7 liver cancer patients (21). Hepatocytes and cholangiocytes were derived from tumor-adjacent normal liver tissues of HCC and CCA patients. TAMs, tumor-associated macrophages; TECs, tumor-associated endothelial cells; CAFs, cancer-associated fibroblasts. **D** H&E and RNAscope staining of *HKDC1*, *HK1*, and *HK2* on the serial sections from the indicated normal human liver and HCC and CCA patient tumors. Solid arrows: normal cholangiocytes; open arrows: normal hepatocytes. Inserts: boxed area in the same image. RNAscope images on the same column share the same scale bar.

We validated the scRNA-seq analysis results using RNAscope on normal human liver and tissues collected from the tumor margin of HCC and CCA patients. The results were consistent with the scRNA-seq analysis (**Figure 2D**). All three HKs were weakly positive in cholangiocytes in the normal human liver, with *HKDC1* showing relatively stronger signals (**Figure 2D, d, i, n**). In HCC, upregulation of *HKDC1* was evident in the tumor cells but not that of *HK1* or *HK2* (**Figure 2D, e, j, o**). In CCA, both *HKDC1* and *HK1* showed significant upregulation, with *HKDC1* being highly specific to tumor cells and *HK1* widespread (**Figure 2D, g & l**). In tumor-adjacent liver tissues of HCC and CCA, *HKDC1* expression remained in cholangiocytes in the portal traids but *HK1* was primarily found in immune cells accumulated in this region (**Figure 2D, f, h, k, m**). *HK2* upregulation was minimal in the HCC and CCA tumors examined, consistent with the scRNA-seq results (**Figure 2D, o-r**). Overall, our data show that HKDC1 is the most tumor cell-specific HK in primary liver cancer.

### HKDC1, HK1 and HK2 are involved in different metabolic activities in HCC cells

Upon the identification of their differential distribution in liver cancer patient tumors, we compared the metabolic activities of HKDC1, HK1, and HK2 in liver cancer cells. Due to the lack of commercial resources for human CCA cell lines, we focused on four human hepatocellular cell lines, including the hepatoblastoma cell line HepG2 and three HCC cell lines, Huh7, PLC/PRF/5, and Hep3B. We found that Huh7 cells showed the lowest endogenous levels of all three HKs (**Figure 3A**), making this cell line as a good model for comparing metabolic changes associated with their ectopic expression. We overexpressed FLAG-tagged *HK1*, *HK2*, and *HKDC1*, individually, in Huh7 cells and confirmed their comparable protein levels (**Figure 3B**). While their individual overexpression led to minor fluctuations in the endogenous levels of the other HKs, these changes were neglectable compared the targeted ectopic expression (**Supplemental Figure 2**). All three HKs promoted Huh7 cell growth in vitro (**Figure 3C**). We then performed a global metabolomic analysis of the HK-overexpressing (OE) Huh7 cells which revealed a distinct metabolic profile in HKDC1-OE cells compared to both the parental control, HK1-OE and HK2-OE cells (**Figure 3D** and **Supplemental Table 1**). HK2-OE caused the least metabolic changes among the three HKs under our experimental conditions. For HKDC1 and HK1, despite their structural similarity, metabolic alterations associated with their overexpression were markedly different (**Figure 3E**). Metabolic pathway enrichment analysis identified a significant impact of HKDC1-OE on the tricarboxylic acid (TCA) cycle (**Figure 3F**). In contrast, HK1-OE substantially upregulated metabolites involved in purine and pyrimidine metabolism pathways (**Figure 3G**). Consistently, intermediate metabolites in the TCA cycle, such as malate, were significantly upregulated in HKDC1-OE cells, whereas key metabolites of purine and pyrimidine biosynthesis pathways, including ribulose 5-phosphate (R5P), inosine and cytidine monophosphate (CMP), were specifically elevated in HK1-OE cells (**Figure 3H**). These findings demonstrate that HKDC1 is functionally non-redundant among three HKs, with HKDC1 being specifically involved in enhancing mitochondrial activity including the TCA cycle to promote tumor growth.

**Figure 3.**
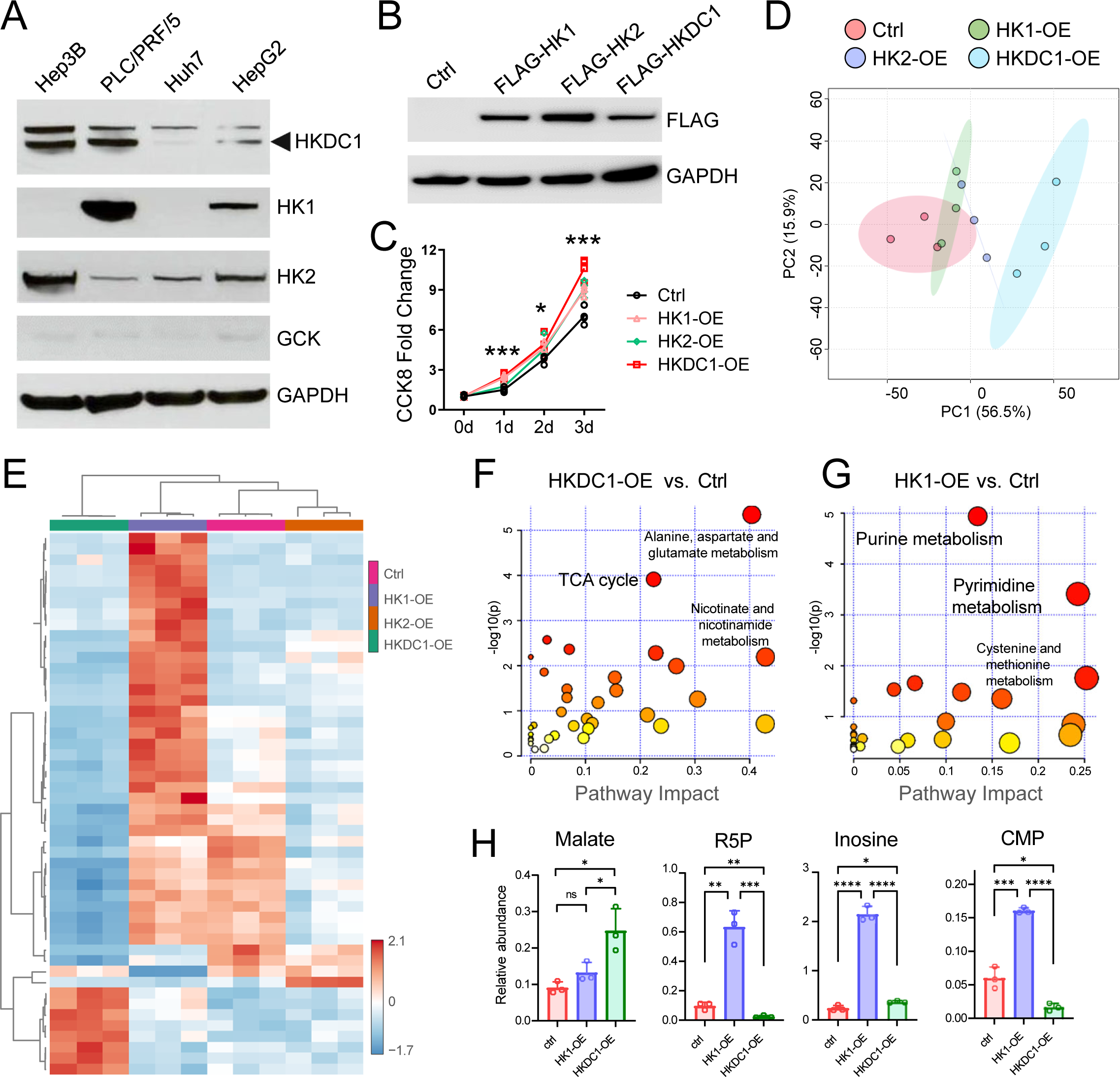
HKDC1, HK1 and HK2 are involved in different metabolic activities in HCC cells. **A** Immunoblotting of the HKs in the indicated hepatocellular cancer cell lines. **B** Immunoblotting of the FLAG-tagged HKs in Huh7 cells. **C** Cell proliferation assay of the HK-OE Huh7 cells. Student *t* test. *P* values: * < 0.05; *** < 0.001. **D** Principal-component analysis (PCA) of metabolomics from Huh7 isogenic cell lines with ectopic overexpression of control, HK1, HK2 and HKDC1, respectively. **E** Heatmap of the significantly altered metabolites among the four isogenic cell lines selected via One-way ANOVA test with *P* values ≤ 0.05. **F** Pathway enrichment analysis of the metabolites differentially upregulated by HKDC1-OE. **G** Pathway enrichment analysis of the metabolites differentially upregulated by HK1-OE. **H** Comparison of the relative abundances of selected metabolites determined by metabolomics in the indicated cell lines. *P* values: * < 0.05; ** < 0.01; *** < 0.001; **** < 0.0001.

### HKDC1 promotes HCC metastasis

Since Hkdc1 was a top upregulated gene in metastatic tumors and organoids in our PPTR and PNR models, we examined its prometastatic functions in the human hepatocellular cell lines. Because of the low levels of endogenous HKDC1 in HepG2 and Huh7 cells (**Figure 3A**), we used these two cell lines to overexpress HKDC1 using an HKDC1-IRES-tdTomato construct (**Figure 4A**). Significantly accelerated cell growth was observed in both HKDC1-OE HepG2 and Huh7 cells (**Figure 4B**), along with their enhanced migration ability (**Figure 4C** and **Supplemental Figure 3**). The protumorigenic function of HKDC1 in liver cancer has been studied in the past, however, only in subcutaneous models (22). Considering the physiological impact of the host liver microenvironment on primary liver cancer, we established orthotopic cell line-derived xenografts (CDXs) by injecting the control and HKDC1-OE HepG2 and Huh7 cells into the liver of NOD scid gamma (NSG) mice (n=5/group). All mice were allowed to reach their humane endpoint to assess their survival. For HepG2, we found no differences in animal survival between the control and OE cells when 1×10^5^ cells/mouse were injected. After slowing tumor development with 1×10^4^ cells/mouse, mice injected with OE cells showed a significantly shorter median survival compared to that controls (**Figure 4D**). Additionally, the OE group developed small intrahepatic metastases in 2 out of the 5 mice but not the control group (**Supplemental Figure 4**). No distant lung metastases were found in either group. For Huh7 cells, there were no differences in animal survival between the control and OE cells regardless of the injected cell numbers (**Figure 4E**). No liver or lung metastases were detected in either group. Upon further tumor examination, we noticed that those generated by HepG2 and Huh7 control cells, despite their low *HKDC1* expression in vitro, showed patchy and high expression of *HKDC1*. Tumors developed from the OE cells showed widespread *HKDC1* expression as expected (**Supplemental Figure 5**). This suggests that HKDC1 expression can be induced in vivo which may explain the limited differences observed in tumor development between the control and HKDC1-OE cells.

**Figure 4.**
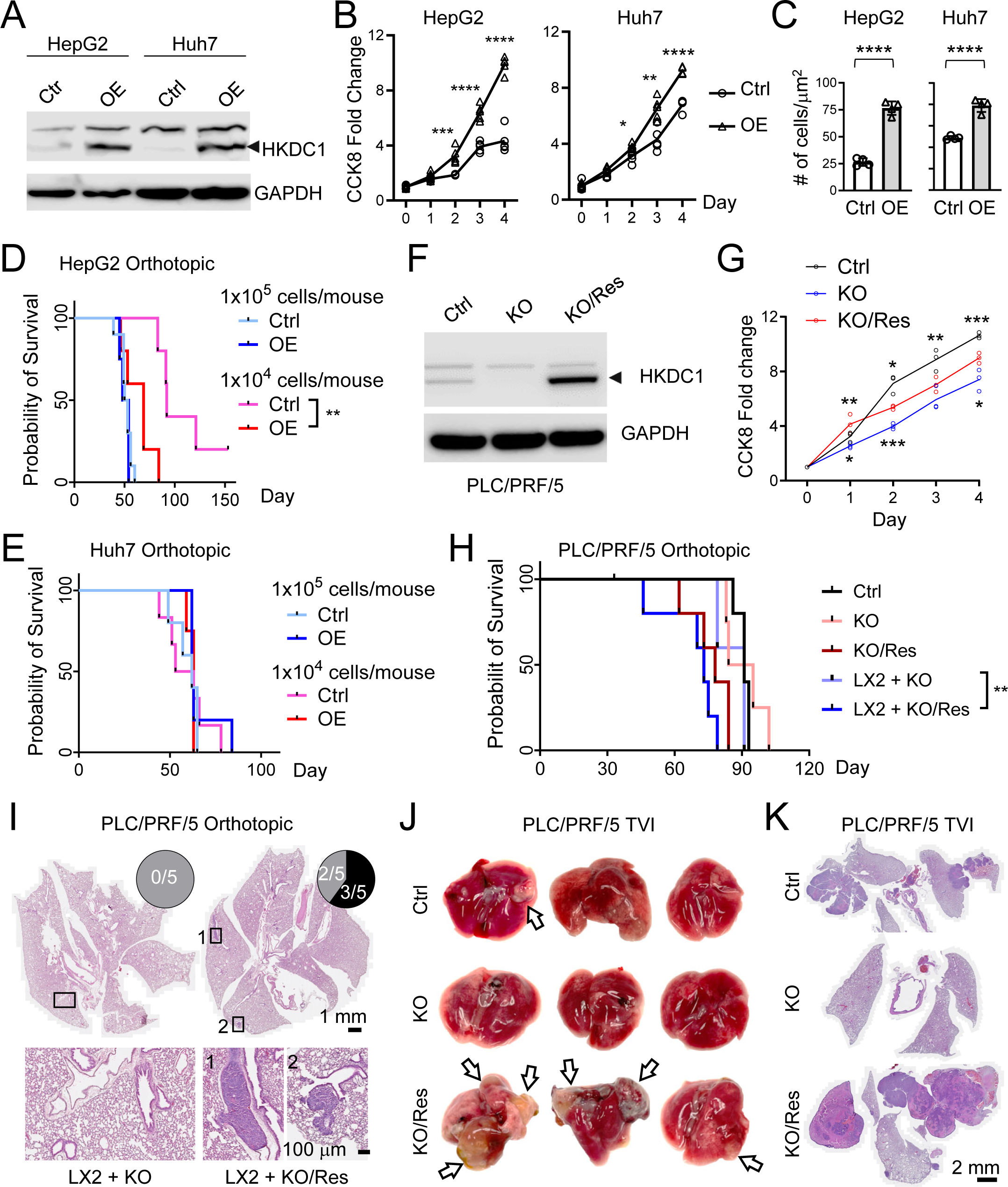
HKDC1 promotes HCC metastasis. **A** Immunoblotting of HKDC1 in the control and HKDC1-OE HepG2 and Huh7 cells. **B** Cell proliferation assay of the control and HKDC1-OE HepG2 and Huh7 cells. **C** Quantification of cell migration assay of the control and HKDC1-OE HepG2 and Huh7 cells. **D** Kaplan–Meier survival curves of mice injected with the indicated HepG2 cells. **E** Kaplan–Meier survival curves of mice injected with the indicated Huh7 cells. **F** Immunoblotting of HKDC1 in the control, HKDC1-OK and -OK/Res PLC/PRF/5 cells. **G** CCK8-based cell proliferation assay of the control, HKDC1-OK and -OK/Res PLC/PRF/5 cells. *P* values on the top of the curves: KO/Res vs. KO; *P* values on the top of the curves: KO vs. control. **H** Kaplan–Meier survival curves of mice injected with the indicated PLC/PRF/5 cells. **I** Top: whole-lung H&E images of mice injected with LX2 mixed with the HKDC1-OK and -OK/Res PLC/PRF/5 cells, respectively; bottom: high-magnification images of the boxed areas in the top image. Pie chart: The number of mice with lung metastasis (black) and with no lung metastasis (grey). Images on the same row share the same scale bar. **J** Gross lung images of the TVI mice injected with the indicated PLC/PRF/5 cells. Arrows: lung metastasis. **K** H&E images of the lung tissues in **J**. Images share the same scale bar. Statistics in **B**, **C**, **G**: Student *t* test. *P* values: * < 0.05; ** < 0.01; *** < 0.001; **** < 0.0001. Statistics in **D** and **H**: Log-rank (Mantel-Cox) test. *P* values: ** < 0.01.

To exclude the potential interference of in vivo *HKDC1* regulation, we generated *HKDC1* knockout (KO) via CRISPR/Cas9 in PLC/PRF/5 cells, a cell line with relatively higher HKDC1 levels among the four cell lines. We subsequently generated *HKDC1*-rescued cells by re-expressing *HKDC1* in the KO cells (KO/Res) (**Figure 4F**). As expected, *HKDC1*-KO PLC/PRF/5 cells showed slowed cell growth in vitro compared to the control cells, and the re-expression of HKDC1 in the KO cells accelerated their growth (**Figure 4G**). However, no statistical differences in survival were observed among mice orthotopically transplanted with the control, KO, and KO/Res PLC/PRF/5 cells (n=5/group) (**Figure 4H**). Given that CAFs are known to promote liver cancer progression (23), we tested co-injection of a human hepatic stellate cell (HSC) line LX2 with the KO or KO/Res PLC/PRF/5 cells as HSCs are the main source of liver cancer CAFs (24). Mice orthotopically injected with LX2 and KO/Res cells showed significantly shorter survival than those injected with LX2 and KO cells (n=5/group) (**Figure 4I**). Moreover, 3 out of the 5 mice injected with LX2 and KO/Res cells developed distant lung metastases while no lung metastases were found in mice injected with LX2 and KO cells (**Figure 4J**). Finally, we tested *HKDC1*-manipulated PLC/PRF/5 cells in a tail veil injection (TVI) lung metastasis model (1×10^6^ cells/mouse, n=3/group). Unlike the limited differences we observed in the orthotopic CDXs, development of lung metastasis was drastically accelerated in mice injected with KO/Res cells compared to KO cells, with the former having significantly more and larger metastases when all mice were examined three months post TVI (**Figure 4K**). Based on these results, we conclude that HKDC1 promotes HCC metastasis although not a particularly strong player. Its prometastatic function may predominantly involve in supporting metastatic growth after tumor cells arrive at a premetastatic niche rather than promoting tumor cell dissemination from the primary site in the liver.

### HKDC1 supports HCC cell growth under hypoxia

Because of the phenotypic differences of HKDC1-manipulated PLC/PRF/5 cells in vitro and in vivo, we compared their transcriptomes via RNA-seq. Gene set enrichment analysis found hypoxia as one of the top enriched pathways when comparing KO vs. control cells and KO/Res cells vs. KO cells (**Figure 5A**). This was not unexpected as cells often undergo metabolic adaption toward enhanced glycolysis under hypoxia (25). Consistently, we also detected a significant and positive correlation between the expression of *HKDC1* and *HIF1A* (**Figure 5B**), a master regulator of hypoxia (26), in HCC and CCA patient tumor RNA-seq profiles available in The Cancer Genome Atlas (TCGA) Program. This correlation was also detected in the PPTR tumors and organoids (**Figure 5C**). RNAscope staining confirmed the colocalization of these two genes in HCC patient tumors and PPTR tumors (**Figure 5D**). Direct evidence supporting the association between HKDC1 and hypoxia came from our observation that cultivation under hypoxia (1% O_2_) for 24 hours induced HIF1α and HKDC1 proteins in all four hepatocellular cancer cell lines we examined (**Figure 5E**). This finding promoted us to examine the impact of HKDC1 loss under hypoxia. We compared the control and two HKDC1-KO PLC/PRF/5 single-cell clones cultivated under normoxia (21% O_2_) and hypoxia. As expected, the KO cells were able to expand although more slowly than the control cells under normoxia. However, under hypoxia, we noticed a complete growth halt of the KO cells during the first 24 hours while the control cells were able to expand similarly to the normoxic condition. The KO cells resumed the growth after 24 hours and then expanded at a similar rate as the KO cells under normoxia (**Figure 5F**). These results suggest that HKDC1 can be induced by hypoxia to provide growth support to HCC cells under acute hypoxic stress.

**Figure 5.**
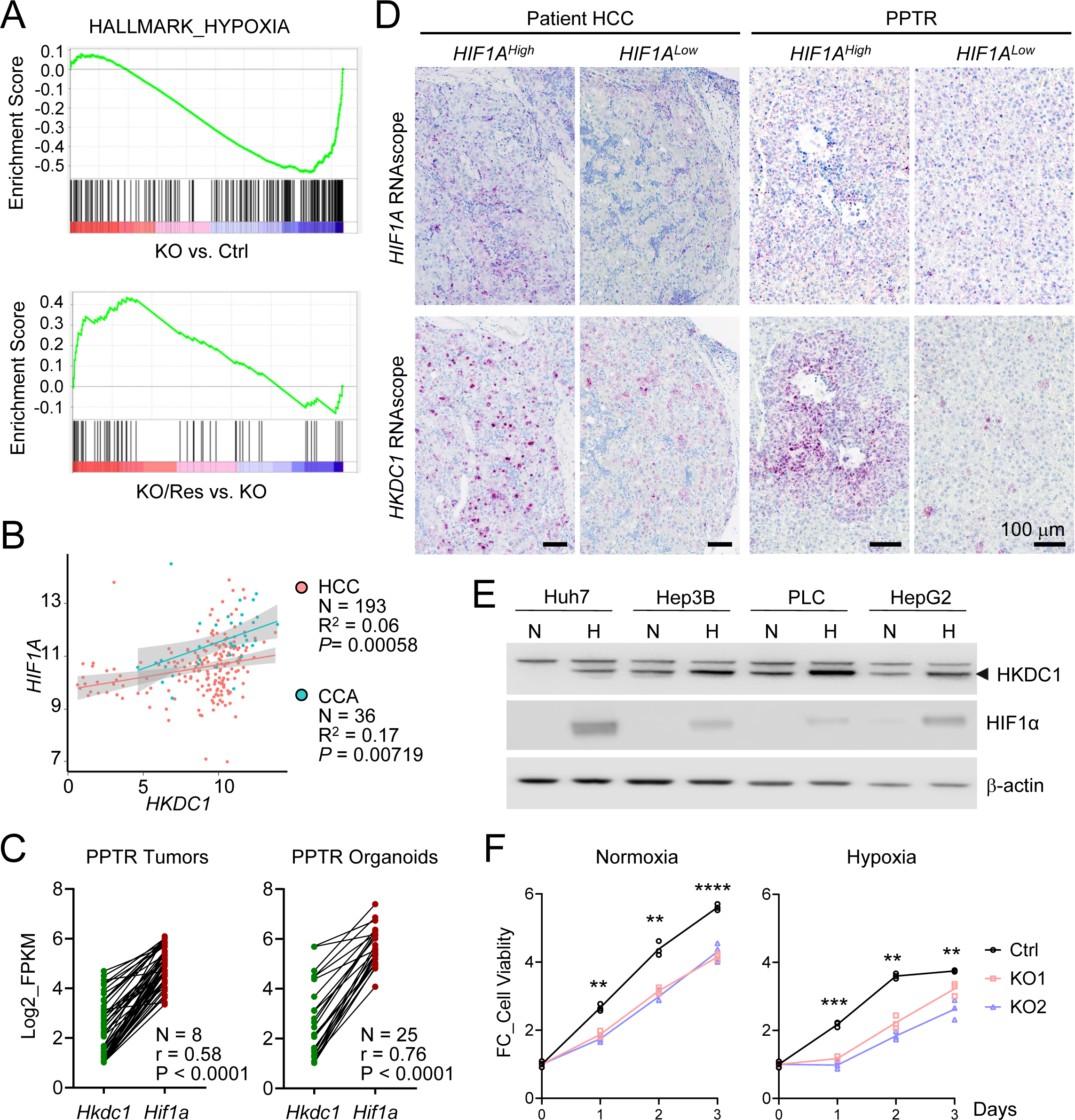
HKDC1 is induced by hypoxia to support HCC cell growth under hypoxia. **A** GSEA analysis found the enrichment of HALLMARK_HYPOXIA gene signature in HKDC1-KO vs. control and KO/Res vs. KO PLC/PRF/5 cells. **B** Spearman correlation analysis of *HKDC1* and *HIF1A* gene expression using RNA-seq data on HCC and CCA patient tumors in TCGA. **C** Spearman correlation analysis of *Hkdc1* and *Hif1a* gene expression in PPTR tumors and organoids. **D** *Hkdc1* and *Hif1a* RNAscope staining on HCC and PPTR tumor serial sections. Images on the same column share the same scale bar. **E** Immunoblotting of HKDC1 in the indicated cells cultivated under normoxia (N, 21% O_2_) and hypoxia (H, 1% O_2_) for 24 hours. **F** Cell proliferation assay of the control and HKDC1-KO PLC/PRF/5 cells under normoxia and hypoxia. *P* values on the top of the curves: control cells, hypoxia vs. normoxia; *P* values on the top of the curves: KO cells, hypoxia vs. normoxia. Statistics in **F** and **G**: student *t* test of the control vs. KO1 cells. *P* value, ** < 0.01; *** < 0.001; **** < 0.0001.

### HKDC1 binds to GSK3**β** to stabilize **β**-catenin and promote liver cancer stemness

The association between HKDC1 and metastasis, organoids, and hypoxia led us to investigate its potential role in stemness properties, given that organoids are known to enrich stem cell population (27, 28), and that hypoxia induces stemness and metastasis in various cancers (29, 30). Indeed, we found that HKDC1-OE HepG2 and Huh7 cells had a significantly higher clonality under the organoid culture condition (**Figure 6A**). Consistently, we observed colocalization of *HKDC1* expression with liver stem cell markers CK19 (31), EpCAM (32), and β-catenin (33, 34) in PPTR genetic tumors, HCC and CCA patient tumors (**Figure 6B**). Molecularly, we studied the potentially interaction between HKDC 1 and β-catenin for the critical role of the β-catenin/WNT signaling pathway in liver cancer stem cells (18, 19). However, no positive association was found between the RNA levels of *Hkdc1* and *Ctnnb1* (the gene encoding β-catenin) in the PPTR and PNR organoids or human HCC and CCA tumors (**Supplemental Figure 6**). In the HKDC1-manipulated PLC/PRF/5 single-cell clones, higher levels of total β-catenin protein were detected in the KO/Res cells while the KO cells showed higher levels of the phosphorylated β-catenin proteins (Ser33/37/Thr41) (**Figure 6C**). Consistently, elevated levels of cyclin D1, a downstream effector of β-catenin (35), were found in the KO/Res cells than the KO cells (**Figure 6C**). Phosphorylation of β-catenin is a key mechanism mediating its ubiquitination and subsequent proteasomal degradation, a process mediated mostly by glycogen synthase kinase 3β (GSK3β) (36). Therefore, we examined the potential interaction between GSK3β and HKDC1. In Huh7 cells overexpressing FLAG-tagged HKDC1, we pulled down GSK3β with the FLAG antibody via immunoprecipitation (**Figure 6D**). Similarly, in PLC/PRF/5 control cells, endogenous HKDC1 was co-immunoprecipitated with GSK3β protein (**Figure 6E**). The binding was less strong compared to the FLAG-HKDC1-OE cells likely due to the much lower level of endogenous HKDC1. When HKDC1-KO PLC/PRF/5 cells showed significantly reduced clonality compared to the control cells, treatment with a GSK3β inhibitor, laduviglusib (also known as CHIR99021) (37), restored the clonality of the KO cells to the levels similar to the KO/Res cells under both the normoxic and hypoxic culture conditions (**Figure 6F and 6G**). We also noticed that colony formation of the KO/Res cells were more robust under hypoxia than normoxia. But laduviglusib treatment did not show difference between these two conditions (**Figure 6F and 6G**). This suggests HKDC1 may have other mechanisms to promote HCC cell stemness under hypoxia besides stabilizing β-catenin. Overall, these results suggest that HKDC1 can bind to GSK3β to stabilizes β-catenin and promotes the stemness properties of HCC cells.

**Figure 6.**
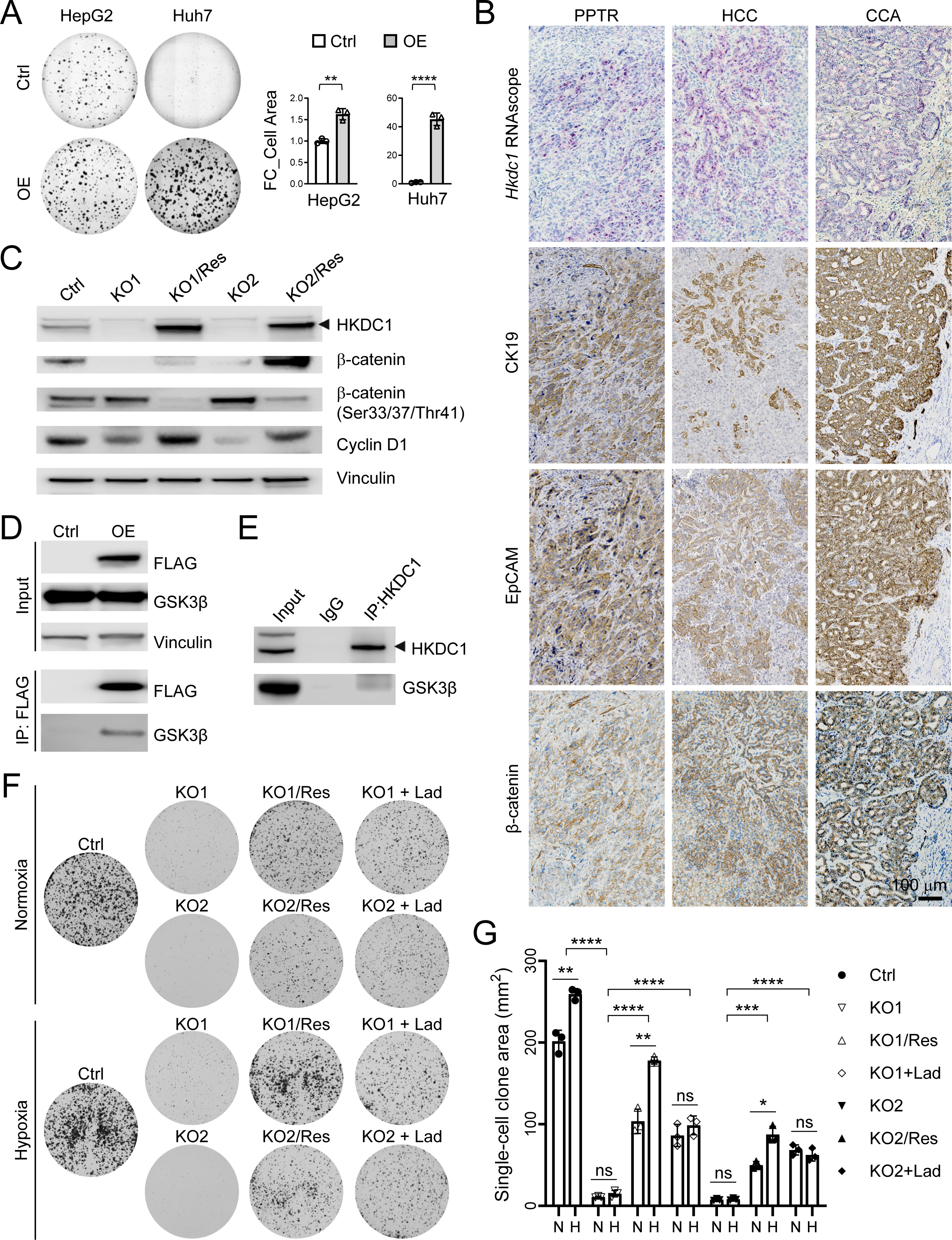
HKDC1 promotes HCC stemness by binding to GSK3**β** and stabilizing **β**-catenin. **A** Organoid stem cell culture of HKDC1-OE HepG2 and Huh7 cells and quantification. **B** RNAscope staining of *Hkdc1* and IHC staining of CK19, EpCAM, and β-catenin on serial sections of tumors from PPTR genetic model and HCC and CCA patients. All images share the same scale bar. **C** Immunoblotting of the indicated proteins in the control, HKDC1-OK and -OK/Res PLC/PRF/5 cells. **D** IP assay of the FLAG in HKDC1-FLAG-OE Huh7 cells pulled down GSK3β protein. **E** IP assay of the endogenous HKDC1 in PLC/PRF/5 cells pulled down GSK3β protein. **F** Colony formation assay of the indicated cells cultured under normoxia and hypoxia. **G** Image-based quantification of the areas of the colonies in **F**. Statistics in **A** and **G**: Student *t* test. *P* values: ** < 0.01; *** < 0.001; **** < 0.0001.

## DISCUSSION

In this study, we identified HKDC1 as a top upregulated gene in the metastatic tumors and organoids from two liver cancer mouse models previously established by our group. We demonstrated that, compared to HK1 and HK2, HKDC1 is upregulated most specifically in the malignant tumor cells in HCC and CCA patient tumors. Our unbiased metabolomic profiling suggests that these three HKs are involved in distinct metabolic activities. Under the experimental conditions we used, HKDC1 is uniquely involved in regulating the TCA cycle in HCC cells. Using HCC orthotopic models, we showed that while HKDC1 can promote lung metastasis although this effect is relatively moderate. However, for HCC cells already arriving in the lung, HKDC1 emerged as a strong driver of their metastatic growth. Finally, we showed that HKDC1 responds to hypoxia, and can promote HCC stemness by binding to GSK3β and stabilizing β-catenin.

HKDC1 is a recently identified HK family member. Our study confirmed its upregulation and protumorigenic roles that have been reported in various cancers including liver cancer (22, 38-43). However, this study is the first to demonstrate its prometastatic function using orthotopic HCC transplantation models. This is also the first study demonstrating the cell type specificity of HKDC1 in the normal and malignant liver in comparison to HK1 and HK2 which are also commonly upregulated in liver cancer. In the normal liver, we found that all three HKs are lowly expressed in cholangiocytes but not in hepatocytes. However, in HCC and CCA patient tumors, HKDC1 expression is specifically upregulated in malignant tumor cells while HK1 and HK2 upregulation is seen across many TME lineages. We believe this finding may provide an explanation to the seemingly contradictory observations published previously that the higher HKDC1 level is associated with poorer prognosis of HCC patients (38) but better survival of CCA patients (43). CCA is well known for its low tumor cellularity due to the abundant TME components (44). More advanced CCA have a greater non-malignant TME compartment which can result in lower tumor cell content and lower HKDC1 levels in bulk tumor-based analyses. Indeed, tumor cells in CCA patient we have examined all show high levels of HKDC1 expression. This highlights the importance to dissect the cellular specificity when it comes to studying molecular drivers of tumors with complex TME such as liver cancer.

Our metabolomic profiling of HK-OE Huh7 cells suggests that these three HKs are involved in distinct metabolic activities. Specifically, HKDC1 demonstrates a pronounced impact on the TCA cycle, while HK1-OE predominantly affects nucleotide metabolism and HK2-OE caused the least metabolic changes in Huh7 cells. This involvement of HKDC1 in TCA cycle is consistent with two recent studies which have revealed that HKDC1 interacts with mitochondria, the organelle where TCA cycle occurs, to support liver cancer progression (22) and prevent cellular senescence (45). Additionally, since mitochondria plays an important role in metastasis by modulating metabolic and genetic responses to dynamic TME cues (46), it supports our observation that HKDC1 responds to hypoxia and needs assistance from CAFs to promote HCC metastasis. CAFs, one of the most critical TME components of liver cancer, are widely known for their roles in creating hypoxic TME and promoting tumor cell dissemination (47, 48). As HKDC1 is known to have a low HK activity and low expression in the normal liver (9, 10), it is conceivable that HKDC1 is not a primary nor classical HK in tumor cells but rather a stress responder, such as under acute hypoxia. It likely collaborates with mitochondria and possibly other organelles and signaling pathways, such as β-catenin/WNT pathway as our study has shown, to maintain energy production in tumor cells under stress. β-catenin/WNT pathway, indeed, is well known for its involvement in glycolysis and metabolic reprogramming (49, 50). This provides an interesting future direction for exploring the mechanistic link between HKDC1 and liver cancer progression from the perspective of tumor-TME interaction. Despite its low expression in normal tissues, the specific upregulation of HKDC1 in tumor cells and metastases makes it a promising therapeutic target in advanced liver cancer, particularly those that have developed distant metastases. Further investigation into the precise roles and regulatory mechanisms of HKDC1 in liver tumorigenesis and metastasis is needed to fully exploit its therapeutic potential.

## MATERIALS AND METHODS

### Mice and Tail vein injection

Animal protocols were approved by the St. Jude Animal Care and Use Committee. All mice were maintained in the Animal Resource Center at St. Jude Children’s Research Hospital. Contral and *HKDC1-*manipulated HepG2, Huh7 and PLC/PRF/5 cells were surgically injected into the liver of two-month-old NSG (NOD scid gamma) mice (The Jackson Laboratory, Bar Harbor, Maine, USA) following the procedure previous reported (17). Kaplan–Meier animal survival curves were graphed using GraphPad Prism 10. *HKDC1-*manipulated PLC/PRF/5 cells were injected into the NSG mice via the tail vein at 1×10^6^ cells/mouse in 100 µl PBS. All mice were maintained in the Animal Resource Center at St. Jude Children’s Research Hospital. Mice were housed in ventilated, temperature- and humidity-controlled cages under a 12-h light/12-h dark cycle and given a standard diet and water *ad libitum*. Animal protocols were approved by the St. Jude Animal Care and Use Committee, and the care and use of experimental animals complied with the animal welfare laws, guidelines and policies.

### Liver cancer patient samples

Deidentified HCC and CCA patient samples were obtained under a protocol approved by the Institutional Review Boards at St. Jude Children’s Research Hospital and The University of Tennessee.

### Cell lines and culture

Human hepatoblastoma cell line HepG2, HCC lines PLC/PRF/5 and Hep3B were purchased from the American Type Culture Collection (ATCC, Manassas, Virginia, USA). HCC cell line Huh7 was a gift from Dr. Jun Yang at St. Jude Children’s Research Hospital. All above cell lines were cultured in Dulbecco’s modified Eagle’s medium (DMEM) containing 1% penicillin-streptomycin (P/S), 1% L-Glutamine, and 10% FBS and incubated in 5% CO2/95% air humidified atmosphere at 37°C. The human immortalized HSC line LX2 was purchased from Millipore Sigma (Cat. No SCC064. St Louis, Missouri, USA) and cultured in DMEM containing 1% P/S, 1% L-Glutamine, and 2% FBS. For cell culture under hypoxia, 1% O_2_ was generated by flushing a 94% N2/5% CO_2_ mixture into the incubator.

### HKDC1, HK1 and HK2 overexpression

pLVX-IRES-tdTomato-3FLAG-HKDC1, pLVX-IRES-tdTomato-3flag-HK1 and pLVX-IRES-tdTomato-3flag-HK2 vectors were generated and packaged into lentiviruses according to the standard procedures. Cells were transduced with lentiviral particles at a M.O.I of 3 and the FASC sorted for tdTomato expression.

### *HKDC1* knockout by CRISPR/Cas9

*HKDC1^KO^*cells were generated using the standard CRISPR/Cas9 technology, sgRNA: 5’ – GGCCCTGGTCAATGACACCGNGG-3’.

### Immunohistochemistry

Immunohistochemistry (IHC) was performed based on the standard protocol. Antibodies used included EpCAM (EPR20532-222, Abcam, Cambridge, MA, USA), β-catenin (#9562, Cell Signaling Technology [CST], Danvers, Massachusetts, USA), CK19 (M0888, Agilent Dako, Santa Clara, California, USA).

### RNAscope staining

RNAscope in situ hybridization of human and mouse *Hkdc1, Hk1,Hk2* and *Hif1a* mRNA transcripts was performed on freshly cut paraffin sections according to the manufacturer’s protocol (Advanced Cell Diagnostics, Newark, California, USA).

### Quantitative RT-PCR

The total RNA in cells were extracted using RNeasy^®^ Mini Kit (#74106, Qiagen, Hilden, Germany) and the concentration and purity of the RNA were measured by nanodrop spectrometer. SuperScript^®^ III First-Strand Synthesis SuperMix for qRT-PCR (#11752-250, Invitrogen, Waltham, Massachusetts, USA) was used for cDNA synthesis from 1 μg total RNA. FastStart Universal SYBR® Green Master (ROX) (#0491385001, Roche, Mannheim, Germany) was used to perform the quantitative PCR assay on the 7900HT Sequence Detection System (Applied Biosystems, Carlsbad, California, USA). The results were analyzed using 2^-ΔΔCt^ Method, With ATCB as the internal reference gene. The primers were as follows: HKDC1: GGCTTCACATTCTCATTTCC (Forward), TGTTGCTGCCTGTTCCTG (Reverse); HK1: GCTCTCCGATGAAACTCTCATAG (Forward), GGACCTTACGAATGTTGGCAA (Reverse); HK2: GATTTCACCAAGCGTGGACT (Forward), CCACACCCACTGTCACTTTG (Reverse); and ATCB: GTTGTCGACGACCAGCG (Forward), GCACAGAGCCTCGCCTT (Reverse).

### Immunoblotting

Cells were lysed using radio-immunoprecipitation assay (RIPA) buffer (#89900, Thermo Fisher Scientific, Waltham, Massachusetts, USA) supplemented with protease and phosphatase inhibitors (#78440, Thermo Fisher Scientific) and 0.5M Ethylenediaminetetraacetic acid (EDTA) (#78440, Thermo Fisher Scientific). Lysates were centrifuged at 14,000 rpm for 15 minutes at 4°C. Protein concentrations were determined using Pierce^TM^ BCA Protein Assay Kit reagent (#23227, Thermo Fisher Scientific) and separated by electrophoresis on NuPAGE^TM^ 4 to 12% Bis-Tris, 1.0, Protein gel (Invitrogen). Antibodies were added according to the manufacturers recommended conditions. Antibodies used include human HKDC1 (NBP182108, Novus Biologicals, Centennial, CO, USA), HK1 (#2024, CST), HK2 (#2867, CST) HIF1α (10006421-1, Cayman Chemistry, Michigan, USA), β-catenin (#9562, CST), β-catenin (Ser33/37/Thr41) (#9561, CST), Cyclin D1 (#2978, CST), GSK3β (#9315, CST), FLAG (F1804, Sigma, St. Louis, Missouri, USA), Vinculin(#13901, CST), β-actin (#4970, CST), GAPDH (#2118, CST).

### Immunoprecipitation

Cells were harvested and lysed with immunoprecipitation buffer (50 mM Tris–HCl, pH 7.6; 150 mM NaCl; 1 mM EDTA; 1% NP-40; 1% protease inhibitor cocktail; 1 mM PMSF). After brief sonication, the lysates were centrifuged at 12000 g for 10 minutes at 4°C. For immunoprecipitation of Flag-tagged proteins, the supernatants were incubated with anti-Flag M2 Affinity Gel (Sigma-Aldrich) at 4°C overnight. Otherwise, supernatants were incubated with indicated antibodies overnight and protein A/G-agarose beads (Santa Cruz, California,USA) for 2 hours at 4°C. The precipitates were washed three times with immunoprecipitation buffer, boiled in sample buffer, and subjected to immunoblot assay.

### Cell proliferation assay

Plates were preincubated in 5% CO2/95% air humidified atmosphere at 37°C and followed by seeding 2 × 10^3^ HepG2 cells, 2000 Huh7 and 2000 PLC cells respectively into per well of 96-well plates. After the cells were cultured for indicated days, 10 μL CCK-8 solutions (CK04, Dojindo Molecular Laboratories, Rockville, Maryland, USA) were added into each well and incubated at 37°C for 2 hours. The absorbance was measured by a Synergy H4 Hybrid Reader (BioTek, Shoreline, Washington, USA) at a wavelength of 450 nm. Cell growth was assessed by cell numbers that had been calculated based on the standard curve between absorbance values and cell numbers. Each sample was triplicate.

### Cell-Titer-Glo cell viability assay

Cells were seeded in a 96 well plate in triplets at 2000 cells/well in 30 ml. Cell Titer Glo (CTG) cell viability assay was conducted daily from Day 1 to Day 7. Briefly, CellTiter-Glo 2.0 (#G9243, Promega, Madison, Wisconsin, USA) was added to each well in a 1:1 v/v ratio. Plates were then covered to keep from light and incubated at RT on an orbital shaker at 150 RPM for 30 mins. After the incubation, plate was read on a synergy H4 plate reader for luminescence.

### Cell migration assay

Cell migration ability was determined by CytoSelect™ 24-Well Cell Haptotaxis Assay (8 µm, Collagen I-Coated, Colorimetric Format; Cell Biolabs Inc, San Diego, CA, USA). Cells suspension were added to the upper chamber (3 × 10^4^ cells in 300 µl blank medium), and 500 µl of media with 10% FBS was added to the lower chamber. After incubation at 37 °C under 5% CO2 for 24 h, the cells on the upper surface were wiped with cotton swabs, while the cells that migrated through the filter pores were stained with cell stain solution for 10 min. The stained cells were observed and photographed under an inverted microscope. Cells were counted from three biological replicates using Image J.

### Organoid growth assay

2 × 10^3^ HepG2 and Huh7 cells were seeded in 48-well plates on matrigel and cultured for 14 days to grow organoids. Areas of the colonies were measured by ImageJ using images from three biological replicates.

### Colony formation assay and laduviglusib treatment

5 × 10^3^ PLC/PRF/5 cells were seeded in 6-well plates and cultured for 7 days with or without laduviglusib (HY-10182, 10 μM, MedChemExpress, New Jersey, USA). Colonies were stained by incubating with 6% Glutaraldehyde (#BP2547-1, Thermo Fisher Scientific) for 20 minutes and 0.05% w/v crystal violet (#C581-25, Thermo Fisher Scientific) for 30 minutes. Plates were imaged on Bio-Rad ChemiDoc Imaging Syster and Image J was used to measure colony area from three biological replicates.

### RNA extraction, sequencing, and data analysis

The total RNA was extracted from cells using RNeasy^®^ Mini Kit (#74106, Qiagen) Kit following the manufacturer’s protocol. Total RNA library was constructed using Illumina TrueSeq stranded mRNA library prep kit and sequenced using the HiSeq 2000/2500 or NovaSeq 6000 platform (2 x 101-bp pair-end reads). On average, we achieved at least 20x coverage for more than 30% of the transcriptome. Gene expression was quantified by STAR(51) (ver. 2.6.0b) under default parameters with the human genome (GRCh37) and annotation file (Gencode v19).(52, 53) Differential expression was performed by ABSSeq under aFold module.(54) Geneset enrichment analysis was performed by GSEA (55).

### Metabolomics analysis by LC-MS/MS

Huh7 cells stably expressing indicated plasmids, were cultured in 6-well plates to ∼85% confluence and washed with 2 mL ice cold 1X Phosphate-Buffered Saline (PBS). Then the cells were harvested in 300 µL freezing 80% acetonitrile (v/v) into 1.5 mL tubes and lysed in the presence of glass beads by Bullet Blender (Next Advance) at 4 °C. Lysed samples were centrifuged at 21,000 × g for 5 min. For each sample, metabolite-containing supernatant was split equally into two aliquots and dried by speedvac respectively. One aliquot was reconstituted in 66% acetonitrile and analyzed by a ZIC_HILIC column (150 × 2.1 mm, EMD Millipore, Burlington, Massachusetts, USA) coupled with a Q Exactive HF Orbitrap MS (Thermo Fisher) in negative ion mode and metabolites were eluted within a 45 min gradient (buffer A: 10mM ammonium acetate in 90% acetonitrile, pH=8; buffer B: 10mM ammonium acetate in 100% H2O, pH=8) (56). Another aliquot was desalted by Ultra-C18 Micro spin columns (Harvard apparatus, Holliston, Massachusetts, USA), resuspended in 5% formic acid and analyzed by acidic pH reverse phase LC-MS/MS with a self-packed column (75 μm × 15 cm with 1.9 µm C18 resin from Dr. Maisch GmbH) coupled with Q Exactive HF Orbitrap MS in positive ion mode. Metabolites were eluted within a 50 min gradient (buffer A: 0.2% formic acid in H2O; buffer B: 0.2% formic acid in acetonitrile). MS settings for both types of samples included MS1 scans (120,000 resolution, 100-1000 m/z, 3 x 10^6^ AGC and 50 ms maximal ion time) and 20 data-dependent MS2 scans (30,000 resolution, 2 x 10^5^ AGC, ∼45 ms maximal ion time, HCD, Stepped NCE (50, 100, 150), and 20 s dynamic exclusion). A quality control sample was injected in the beginning, middle and the end of the samples to monitor the system stability of LC-MS.

The data analysis was performed by in-house software suite JUMPm (57). Raw files were converted to mzXML format followed by peak feature detection for individual sample and feature alignment across samples. Metabolite identification were supported by matching the retention time, mass/charge ratio, and MS/MS fragmentation data to our in-house authentic compound library, downloaded experimental MS/MS library (MoNA, https://mona.fiehnlab.ucdavis.edu/) and mzCloud (https://mzcloud.org). Peak intensities were used for metabolite quantification. The data was normalized by both cell numbers (before data collection) and trimmed median intensity of all features across samples (post data collection) (58). The normalized metabolomics data were applied to the multivariate and statistical analyses. The pathway enrichment analysis of differential metabolites among groups and the heat map with clustering of samples and differential metabolites were performed using Metaboanalyst 6.0 (http://www.metaboanalyst.ca).

### Statistical analysis

Student’s t-test was used to evaluate differences between two groups. One-way ANOVA test was used to compare >=3 groups. A *P* value of less than 0.05 was considered statistically significant.

## DISCLOSURES

The authors have declared that no conflict of interest exists.

## AUTHOR CONTRIBUTIONS

L.F. conducted most of the biological experiments. The order of the co-first authors was assigned based on their intellectual contributions. C.T., H.T., X.L. conducted experiments. W.Y., L.M., and M.N. conducted the computational analyses. J. P., J. Y., and E.S.G. provided technical and intellectual support. L.Z. conceived and oversaw the research. All authors contributed to the writing of the manuscript.

## ABBREVIATIONS

HCC: hepatocellular carcinoma
CCA: cholangiocarcinoma
HK: hexokinase
HKDC1: hexokinase domain containing 1
TME: tumor microenvironment
PPTR: Prom1^CreERT2^; Pten^flx/flx^; Tp53^flx/flx^; Rosa-ZsGreen
CAF: cancer-associated fibroblast
TAM: tumor-associated macrophage
TEC: tumor endothelial cell
TCA: tricarboxylic acid
R5P: ribulose 5-phosphate
CMP: cytidine monophosphate
OE: overexpression
KO: knockout
CDX: cell line-derived xenograft
NSG: NOD scid gamma

## Notes

**FINANCIAL SUPPORT** This work was supported by American Cancer Society Research Scholar Grant RSG-18-026-01 (to L.Z.).

### Competing Interest Statement

The authors have declared no competing interest.

