## Supplemental Figures 1-7 for "HKDC1 Promotes Liver Cancer Stemness Under Hypoxia via Stabilizing β-Catenin"

Supplemental Figure 1

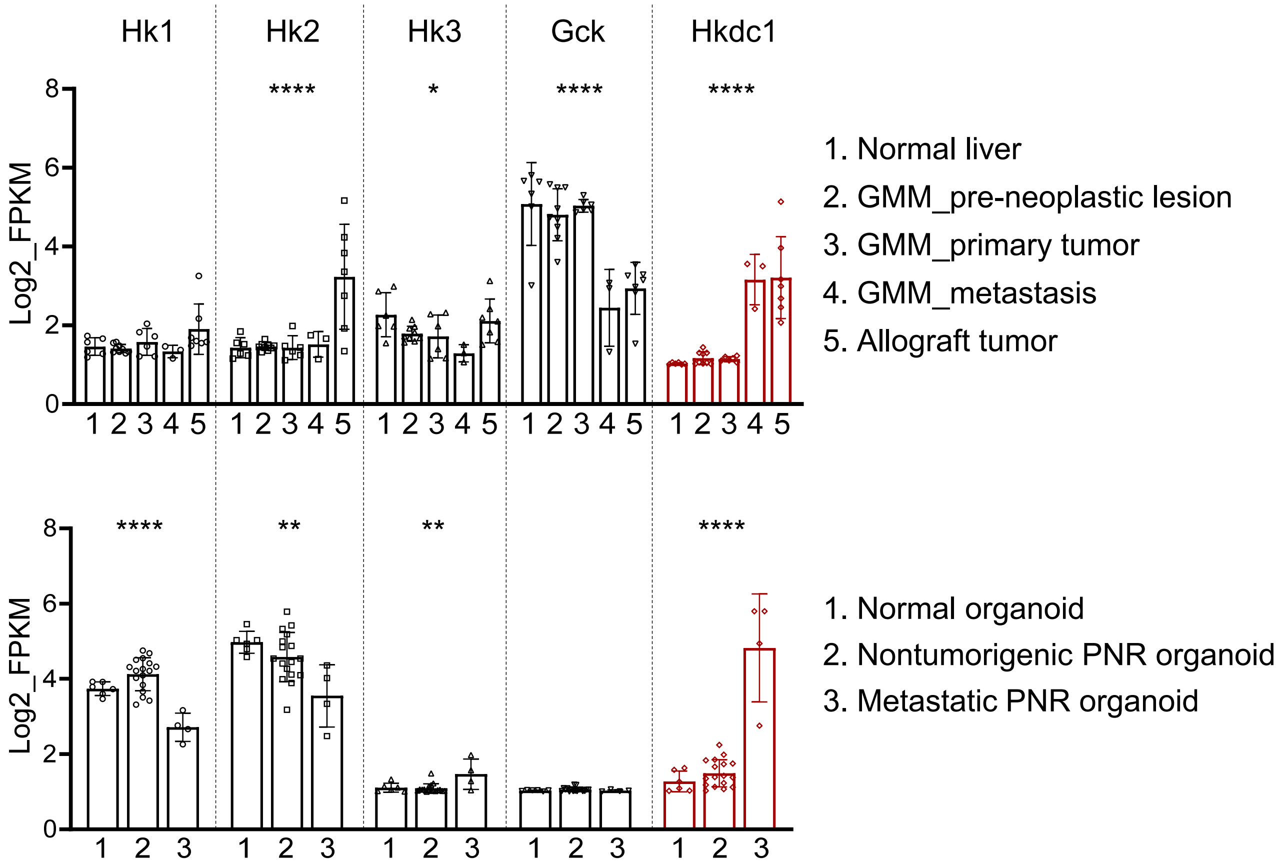


**Supplementary Figure 1. Hkdc1 expression is associated with the tumor progression in the PNR mouse model.**

Quantitative comparison of the expression of five HKs in the PNR tumor tissues (N = 6, 9, 6, 3, and 7, respectively, for the five groups presented) and organoids (N = 6, 17, and 4, respective, for the three groups presented) in the indicated groups. One-way ANOVA test. P values: * < 0.05; ** <0.01; ****<0.0001.

### Supplemental Figure 2

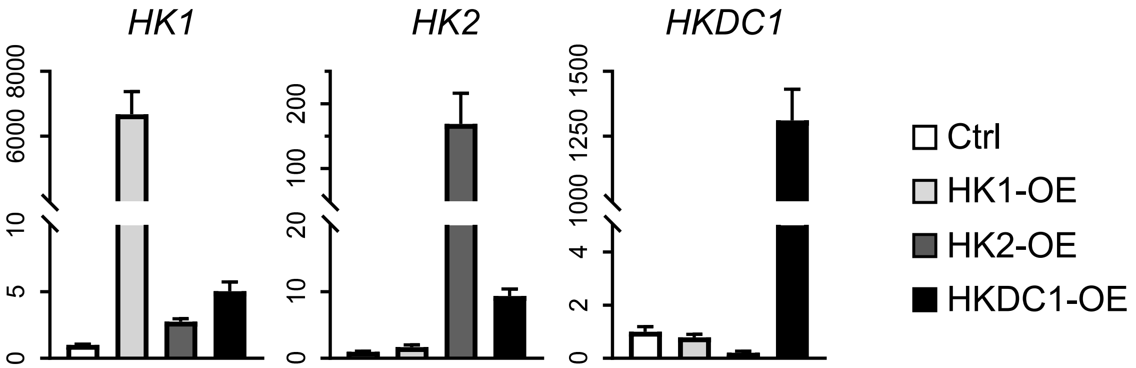


**Supplementary Figure 2. Ectopic expression of the individual HK in Huh7 cells caused minor disturbance in the endogenous HK expression.**

Quantitative RT-PCR of *HK1*, *HK2*, ***and*** *HKDC1* in the control and HK1-, HK2- and HKDC1-OE Huh7 cells.

Supplemental Figure 3

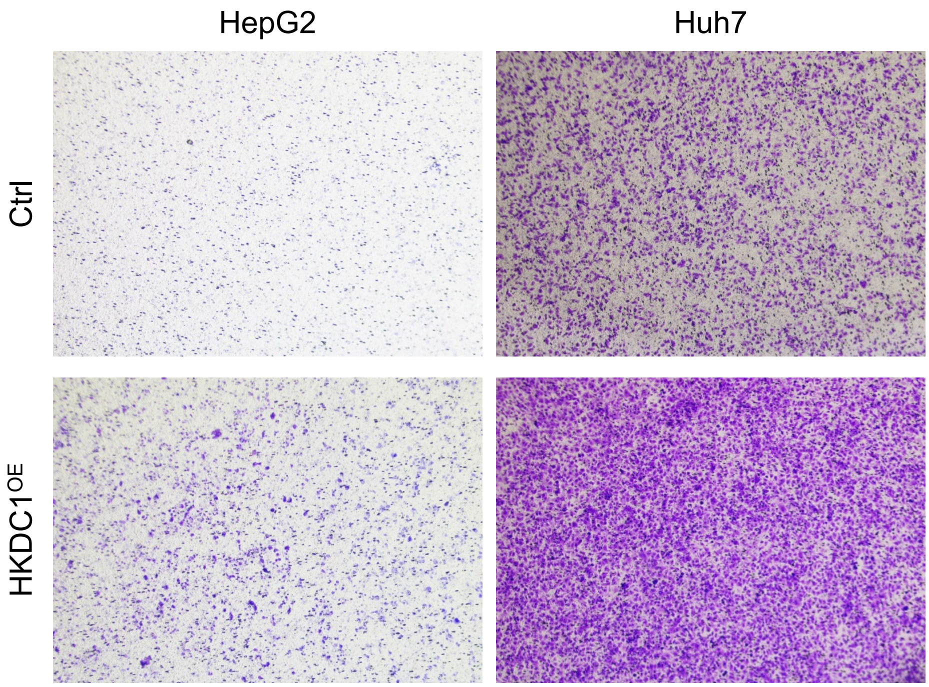


**Supplementary Figure 3. Hkdc1 overexpression promotes HepG2 and Huh7 migration in vitro.**

Crystal violet staining of the indicated cells subjected to 24 hours of the transwell migration assay.

### Supplemental Figure 4

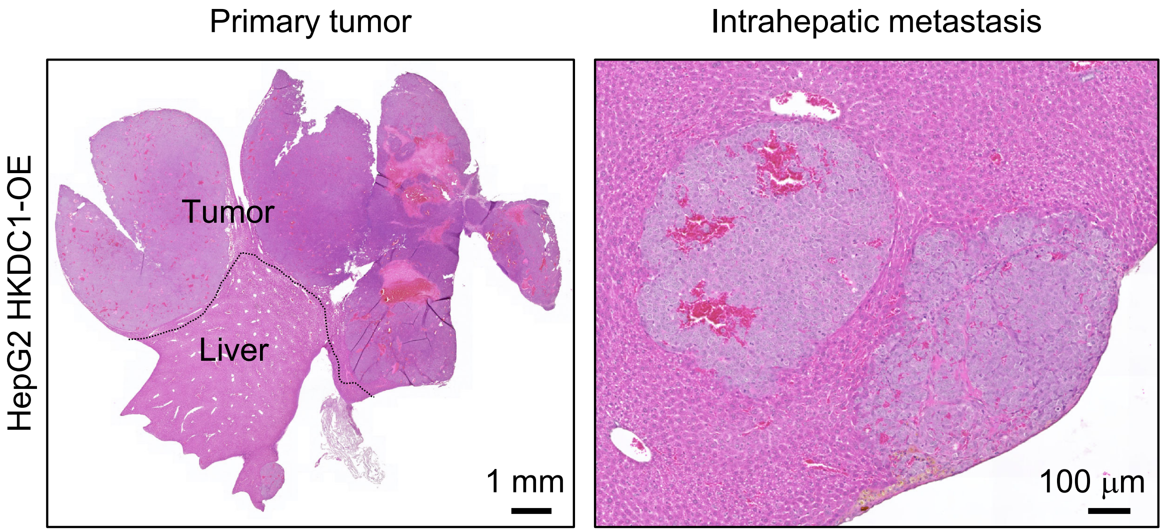


**Supplementary Figure 4. HKDC1-OE HepG2 cells developed intrahepatic metastasis in the orthotopic CDX.**

H&E staining of the liver from a mouse injected with indicated HKDC1-OE HepG2 cells.

### Supplemental Figure 5

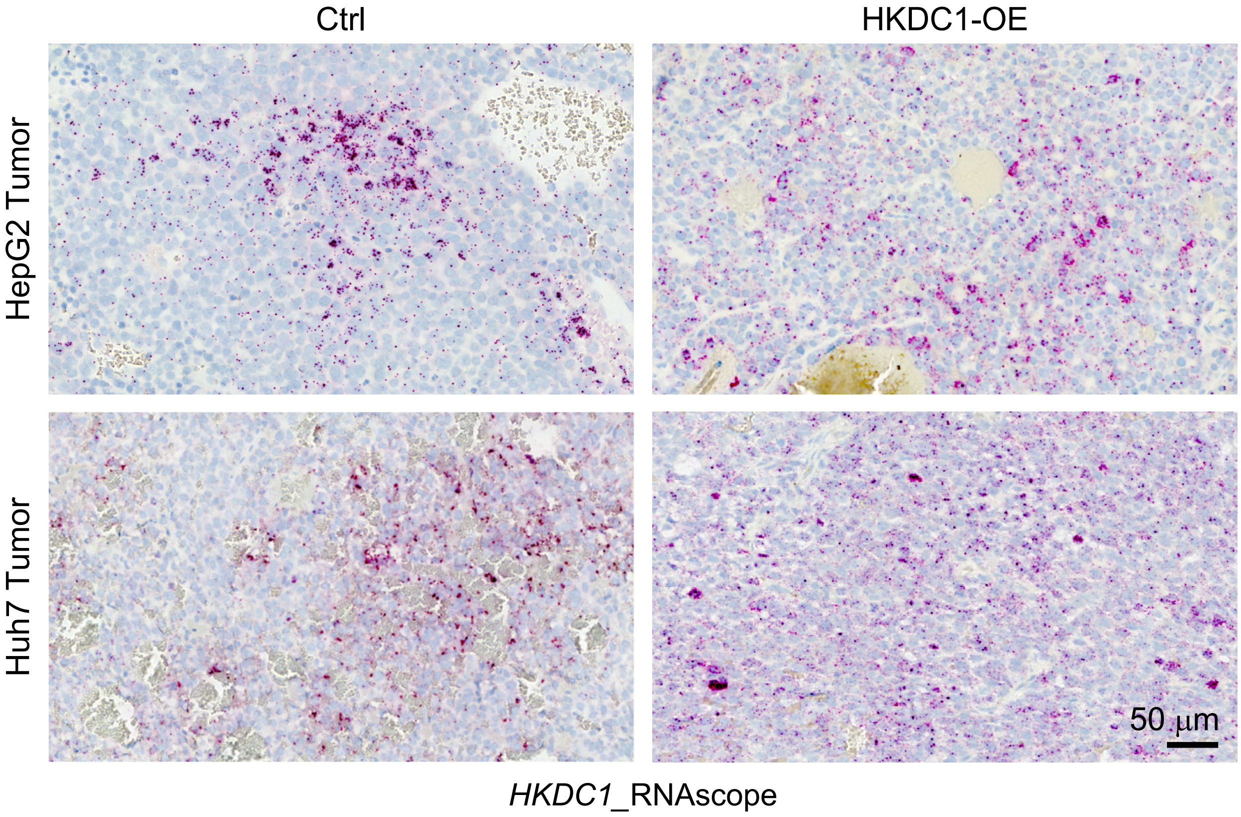


**Supplementary Figure 5. HKDC1 expression in HepG2 and Huh7 orthotopic CDX tumors.**

*HKDC1* RNAscope on the liver tumors generated by the control and HKDC1-OE HepG2 and Huh7 cells.

### Supplemental Figure 6

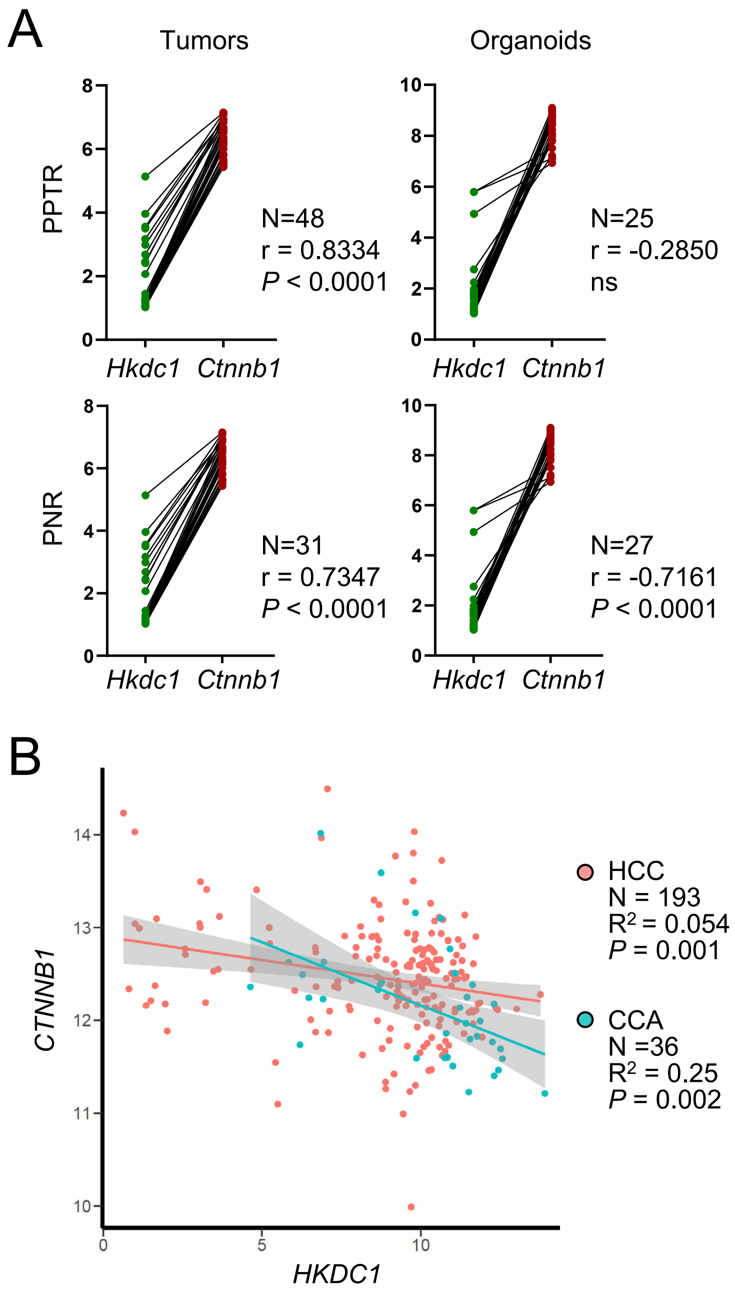


**Supplementary Figure 7. HKDC1 and β-catenin gene expression in mouse and human liver cancer samples.**

**(A)** *Hkdc1* and *Ctnnb1* gene expression are positively correlated in PPTR and PNR tumor tissues but not organoids.

**(B)** *HKDC1* and *CTNNB1* gene expression are not positively correlated in human HCC and CCA tissues.
